# Parietal stimulation reverses age-related decline in exploration, learning, and decision-making

**DOI:** 10.1101/2023.10.21.563408

**Authors:** Eun Jung Hwang, Sayli Korde, Ying Han, Jaydeep Sambangi, Bowen Lian, Ama Owusu-Ofori, Megi Diasamidze, Lea M. Wong, Nadine Pickering, Sam Begin

## Abstract

Aging can compromise decision-making and learning, potentially due to reduced exploratory behaviors crucial for novel problem-solving. We posit that invigorating exploration could mitigate these declines. Supporting this hypothesis, we found that older mice mirrored human aging, displaying less exploration and learning during decision-making, but optogenetic stimulation of their posterior parietal cortex boosted initial exploration, subsequently improving learning. Thus, enhancing exploration-driven learning could be a key to countering cognitive aging.

## MAIN

Experience and learning can enhance the quality of our decisions. However, older adults often face challenges in making adaptive decisions, as their cognitive ability to learn new rules and skills decline^1–6^. This reduced learning ability may be associated with a decreased propensity for exploring novel environments and behaviors^6–8^. When confronted with an unfamiliar task, the early-phase exploration of the environment and behavioral repertoire could facilitate the discovery of a task-appropriate solution^9–11^. Thus, exploration, typically characterized by behavioral variabilities, could influence learning. In keeping with this view, studies have associated faster and more effective learning with a greater variability in task-related behavior before or during the initial training phase of a new task^12,13^. Animal studies have further demonstrated that suppressing exploration impairs spatial and sensorimotor learning, highlighting the necessary role of exploration in the learning process^10,14^.

Growing evidence links exploratory decisions to the posterior parietal cortex (PPC). The PPC is strongly engaged when people make exploratory decisions during reinforcement learning^15^. Inhibitory stimulation of the PPC suppresses visual exploration in the space contralateral to the stimulation^16^. In mice, PPC neurons represent decision biases including exploration that depend on previous trial history, and perturbing PPC activity alters those history-dependent biases^17–19^. These findings suggest that the age-related decline in exploratory decision-making could potentially be counteracted by manipulating PPC activity. Additionally, the correlation between exploration and learning implies the compelling possibility that restoring exploratory behavior may enhance the ability to make learning-dependent decisions.

To investigate this hypothesis, we initially examined age-related alterations in decision making and learning using the standardized international brain laboratory (IBL) decision-making task in mice^20,21^. In this task, mice are presented with a Gabor patch stimulus on either the right or left side of the screen and turn a wheel in the direction that moves the stimulus to the center of the screen to receive a water reward (**Figure 1a**). Specifically, leftward turning for right side stimuli and rightward turning for left side stimuli lead to a reward. Previous studies have shown that young adult mice can learn this stimulus-choice-outcome rule through trial-and-error over multiple sessions^20,21^. However, the impact of age on behavior in the IBL task has not been examined. Therefore, we trained mice in this task for approximately 40 sessions (1 session per day) across four age groups (G1-G4; **Figure 1b**). Consistent with previous findings, we observed a gradual improvement in the correct choice rate across sessions in G1, the youngest group (N=21; 2.7 ± 0.18 (mean ± s.d.) months old) (**Figure 1c**). The majority of G1 mice (20 out of 21 mice; 95%) achieved a correct choice rate greater than 80% in 25 ± 8.5 (mean ± S.D.) days (**Figure 1d**). In contrast, only 2 out of 16 mice in G4, the oldest group (N=16; 20.6 ± 0.60 (mean ± s.d.) months old) achieved a correct choice rate greater than 80% (**Figure 1d**). This difference in the fraction of successfully trained mice between G1 and G4 is statistically significant (Chi-square test, χ^2^(2, N=37) = 37.9, p<0.001). Accordingly, the highest correct choice rate achieved across all sessions decreased with age (**Figure 1e**). We also quantified the learning rate of each mouse by calculating the time constant of an exponential function fitted to their learning curve (Methods; **Figure S1**). The learning rate gradually decreased with age, with a significantly higher learning rate observed in G1 compared to G4 (Wilcoxon rank-sum test, z =, p<0.001) (**Figure 1f**). These findings collectively indicate that the acquisition of a novel decision-making rule slows down with age. Moreover, this age-related decline in learning was consistent across sexes and genotypes (**Figure S2**).

**Figure 1.**
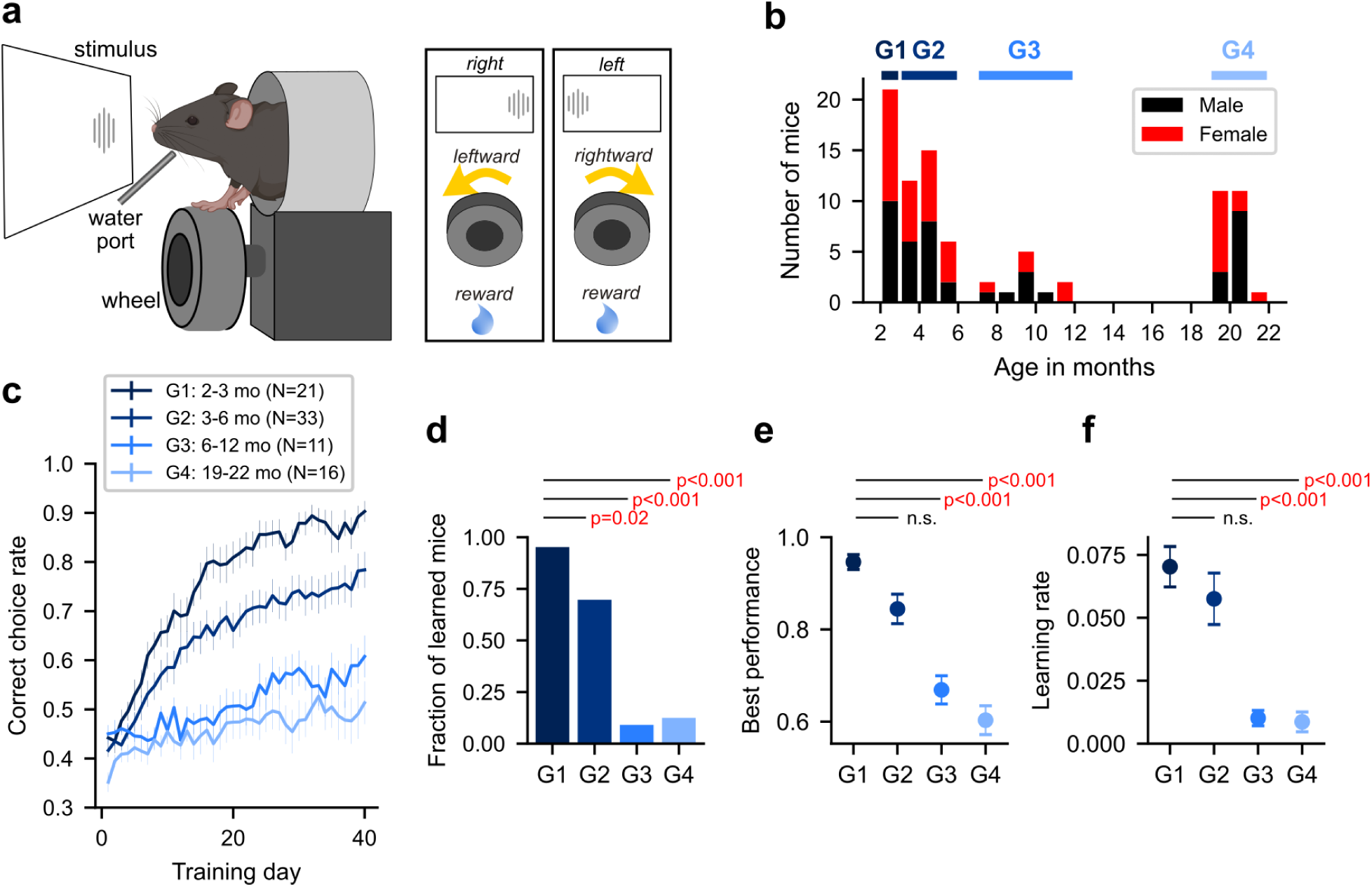
Learning-dependent decision making declines with age. **a.** Decision-making task. (left) international brain laboratory task set-up. (right) decision-making rule related to stimulus, choice, and outcome. **b.** Distribution of age and sex across trained mice. The thick horizontal bars above the histogram indicate the age interval included in each of 4 age groups. **c.** Learning curves from 4 age groups. X-axis represents training day (1 session / day), and y-axis the correct choice rate. Mean ± S.E. **d.** The fraction of learned mice in each group. **e.** The best performance across all training days. Mean ± S.E. **f.** The learning rate. Mean ± S.E.

In contrast to the decision-making rule learning, repetition-based procedural motor learning was largely unaffected and, in some cases, even enhanced by age. We analyzed kinematic metrics of wheel movements during the IBL task, including response time, movement speed, and trial-to-trial correlation of velocity profiles. Consistent with the consensus that processing speed and movement vigor decrease with age, we observed significant differences in baseline response time and movement speed between G1 and G4 at the beginning of training (**Figure S3**). However, all measures of motor learning improved at similar rates across all age groups, and the kinematic metrics at the end of training were not statistically different across age groups (**Figure S3**). Thus, the decreased learning capacity with age is not uniform across all types of learning, and the impaired decision-rule learning does not indicate a general lack of motivation. The neural circuit supporting simple repetition-based sensorimotor learning appears to remain relatively intact at later stages of life^22^.

As humans age, they tend to exhibit less exploratory choices in ambiguous and uncertain decision-making contexts, even if such exploration could lead to more favorable outcomes^7^. For example, in the Iowa gambling task, older adults are less likely to switch their choices after a winning trial and thus slower in finding the most profitable option compared to younger adults^6,8^. We aimed to determine whether mice display this pattern - reduced exploratory choices in ambiguous contexts - as well. In our task, the early training sessions provide such contexts as mice have not acquired the correct decision rule yet. Thus, to quantify the degree of exploratory decisions in an ambiguous context, we first calculated the fraction of switch trials following a reward (’win-switch’) during the first 3 training sessions (**Figure 2a**). Our results showed a significant difference in the fraction of win-switch trials across the age groups (Kruskal-Wallis one way ANOVA, H(3) = 23.7, p<0.001), with a significant reduction in G4 compared to G1 (Wilcoxon rank-sum test, z = 4.5, p<0.001) (**Figure 2b**). This finding indicates that older mice, like humans, exhibit fewer ‘win-switch’ choices and more ‘win-stay’ choices. We also observed that the fraction of ‘stay’ choices increases with age, regardless of the previous trial outcome (**Figure 2b**). This increased tendency to repeat the same choice irrespective of previous choice outcome would also contribute to the decrease in exploratory decisions.

**Figure 2.**
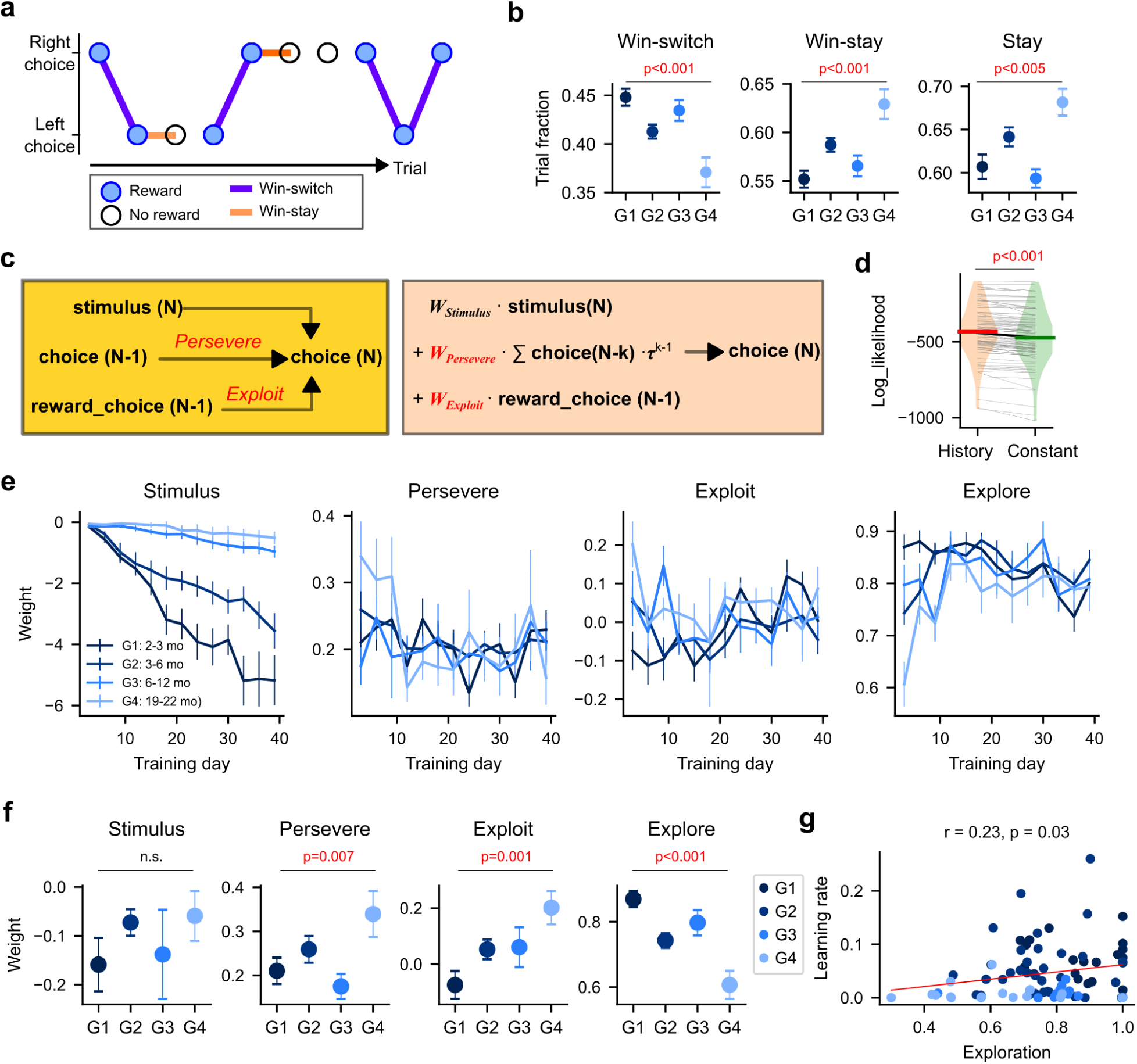
Early exploratory decisions decline with age. **a.** A snippet of choice sequence from a young mouse. The filled circles indicate rewarded trials. Purple lines represent win-switch, while orange lines win-stay. **b.** The fraction of win-switch, lose-switch, and switch during the first 3 sessions. Mean ± S.E. **c.** History-dependent choice model. Choice on trial number N depends on the N-trial stimulus, previous choices (i.e., perseverance), and previous rewarded choice (i.e., exploitation). The model also includes outcome history and constant terms that are not included in the illustration for brevity. **d.** The log-likelihood distribution of the full model with history terms versus a partial model that consists of only stimulus and constant. **e.** The estimated dependence on the current trial stimulus and trial history across learning phases (computed for every, non-overlapping 3 consecutive sessions). Mean ± S.E. **f.** Weights corresponding to the degree of stimulus dependence, perseverance, exploitation, and exploration during the first 3 sessions. Mean ± S.E. **g.** The correlation between learning rate and early exploration. The dots represent individual mice.

To systematically quantify these different trial-history dependent decision strategies, we utilized a behavior model in which choice on a given trial is influenced by the choice and outcome of past trials, in addition to the current trial stimulus information^17^ (Methods; **Figure 2c**). This model includes terms to represent the degrees of ‘stay’ (perseverance) and ‘win-stay’ (exploitation) strategies, respectively. The inverse of these two strategies (i.e., negative weights) would indicate exploratory behavior. We estimated the weight of each term by fitting the model to the choice sequence of each mouse across different phases of learning. The history-dependent model outperformed a simple model that consists of only stimulus and constant in 94% of mice (Likelihood ratio test between the two models; p<0.05) and predicted the choice sequence of each mouse significantly better than the simple model (Wilcoxon signed-rank test, z = 73, p<0.001) (**Figure 2d**). As shown in **Figure 2e**, choice-dependence on stimulus increases with training but at different rates in different age groups, echoing the age-related decline of the learning rate. We also noticed differences in trial-history dependence in early training sessions. For instance during the first 3 sessions, stimulus dependence is nearly zero in all age groups, confirming that mice did not acquire the new decision rule yet (**Figure 2f**). In this novel ambiguous context, however, both perseverance and exploitation increase with age, with each showing a significantly higher value in G4 than in G1 (Wilcoxon rank-sum test, z = 3.9, p<0.001;Wilcoxon rank-sum test, z = 2.1, p=0.03) (**Figure 2f**). Notably, the exploitation term had a negative weight in G1 mice, indicating that the youngest mice suppress the exploitative tendency in favor of an exploratory strategy in the unfamiliar setting. As a comprehensive measure of exploration, we computed an exploration score by combining the degrees of perseverance and exploitative strategy (Methods). The compound exploration score was significantly lower in G4 than in G1 (Wilcoxon rank-sum test, z = 4.0, p<0.001) (**Figure 2f**). These results were comparable across variants of behavior models (**Figure S4**), consistently indicating that older mice exhibit reduced exploratory behavior in the novel context. The difference in exploration gradually disappears in later training sessions, as older mice become less persevering and exploitative, while younger mice become slightly more exploitative and heavily rely on stimulus information (**Figure 2e**).

We then investigated whether the extent of exploration early in training was correlated with subsequent learning of the choice rule. We observed significant correlations between exploration in the initial training sessions and the learning rate (**Figure 2g**). Mice that made more exploratory choices early on displayed faster rule learning. Although this correlation was not significant within each age group, the group-level differences in exploration were sufficiently robust to account for the group-level differences in learning.

The observed correlation between the exploratory score and learning performance could be attributed to shared or correlated inputs to independent circuits, each controlling exploration and learning respectively. For instance, neuromodulatory signals may alter with age, making global impacts on both functional circuits concurrently^23^. Alternatively, the exploration circuit may directly interact with the learning circuit, promoting the acquisition of novel stimulus-choice-outcome rules. The latter hypothesis predicts that activating the exploration circuit in old mice may improve their learning. In our previous studies, we found that inactivating the PPC during the inter-trial interval (ITI) weakens trial history dependence, including exploratory bias in a familiar task^17,18^. In addition, PPC inactivation increases choice perseverance during early learning^24^. These previous findings suggest that manipulating PPC activity can alter exploratory-related behavior. To test our prediction, we trained a new group of old mice (G4-opto; N=7; 19.9 ± 0.05 (mean ± s.d.) months old) in the same decision-making task while stimulating their PPC.

To stimulate the PPC, we expressed an excitatory opsin ChRmine and implanted fiber optic cannulas in the PPC bilaterally (**Figure 3a**)^25^. During the initial 15-16 sessions of the IBL task, we delivered red LED light (625 nm; ∼10 mW) to the PPC during the ITI on approximately 23% of randomly selected trials (**Figure 3b**). We found that exploitation during the first 3 training sessions was significantly lower in G4-opto compared to G4 mice (Wilcoxon rank-sum test, z=2.0, p=0.04; z=2.1, p=0.03) (**Figure 3c**). Consequently, the exploration score was significantly higher in G4-opto than in G4 mice (Wilcoxon rank-sum test, z=0.5, p=0.61) (**Figure 3c**). To examine whether the boost of exploration and suppression of exploitation in the G4-opto group occurred selectively in stimulated trials, we estimated the history dependence separately for LED-on and off trials (**Figure 3e**). Although the general effect size was larger in the LED-on trials, we found significant boost of exploration and suppression of exploitation even in LED-off trials. Therefore, the PPC stimulation effects appear to outlast the stimulated trials, enhancing exploratory choices in general.

**Figure 3.**
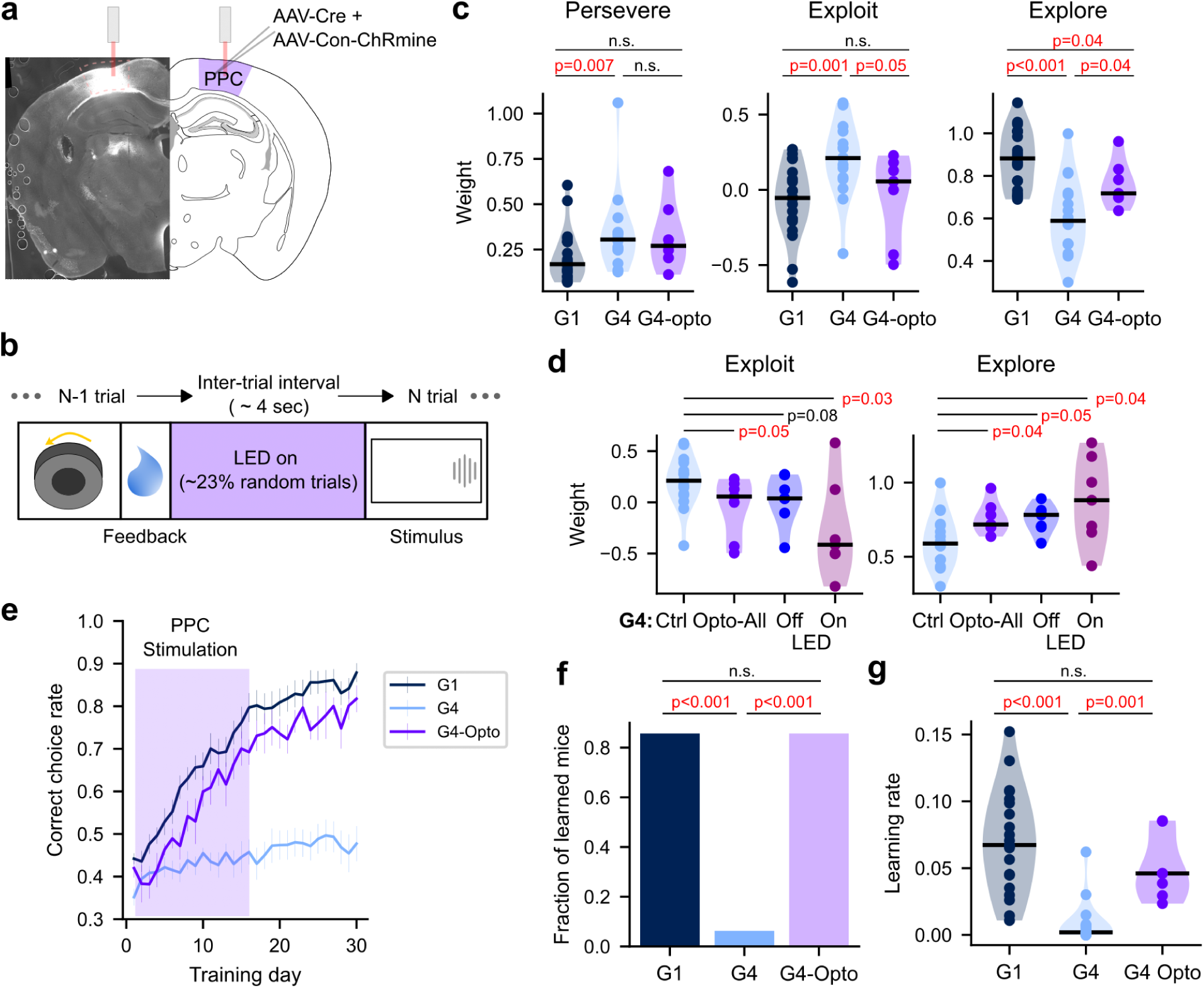
PPC stimulation in aged mice restores exploration and learning. **a.** Chrmine, the red-shifted excitatory opsin, was virally expressed and fiber optic cannulas were implanted in PPC bilaterally. The brain slice on the left demonstrates a successful expression of Chrmine in PPC. **b.** In optogenetic sessions (first 16 training sessions), on randomly selected trials (∼23%), LED light was turned on during the inter-trial interval. **c.** The degree of perseverance, exploitation, and exploration in G1, G4, and G4-opto groups. G1 and G4 are the same as in Figures 1 and 2. G4-opto matches G4 in age, but PPC was stimulated. Dots: individual mice. **d.** The comparison of exploit and exploration across the trial subgroups in G4: all trials from the control group, all trials from the optogenetic group, LED-off trials from the optogenetic groups, and LED-on trials from the optogenetic group. **e.** The learning curves. Mean ± S.E. **f.** The fraction of learned mice within 30 training days. **g.** The learning rate.

Our hypothesis predicted that enhancing exploration in old mice would improve their learning. To test this prediction, we compared learning curves (**Figure 3d**). Remarkably, 6 out of 7 old mice with PPC stimulation reached an over 80% correct choice rate within 30 sessions, which is significantly higher than the fraction observed in unstimulated old mice (Chi-square test, H(2)=14.5, p<0.001) (**Figure 3e**). Furthermore, the learning rate of G4-opto mice was significantly higher than that of unstimulated G4 mice (Wilcoxon rank-sum test, z=3.3, p=0.001) and comparable to that of unstimulated G1 mice (Wilcoxon rank-sum test, z=1.2, p=0.22) (**Figure 3f**). Therefore, PPC stimulation increased exploratory decisions early in training and subsequently enhanced the learning of the decision rule, enabling old mice to achieve a higher rate of correct choices after training. These results support our hypothesis that exploration in the early training phase can promote learning, leading to improved decision-making performance.

The precise mechanisms underlying the PPC-stimulation-induced enhancement of exploration and learning remain unknown. The PPC is a prominent region that displays cortical thinning with age, and the extent of PPC thinning correlates with a reduction in risky, uncertain decisions in humans^26^. Additionally, cortical atrophy in the PPC is associated with impaired associative learning^27^. Given these observations in humans, we conjecture that our optogenetic stimulation might have reversed age-related activity changes in the PPC, reinstaining initial exploratory behavior and subsequently restoring learning and decision-making abilities. Taken together, our results suggest promising strategies to intervene in age-related cognitive decline in humans; non-invasive brain stimulation targeting the PPC or behavioral therapy aimed at promoting exploration-driven flexibility. These approaches could potentially provide assistance to older adults grappling with the acquisition of new decision rules and the navigation of challenging tasks.

## METHODS

### Animals

All procedures were in accordance with protocols approved by the Rosalind Franklin University Institutional Animal Care and Use Committee and guidelines of the National Institute of Health. Five different strains of mice (both sexes) were used; wild type C57Bl/6J [JAX 000664] (24 female, 22 male), B6.Cg-Gt(ROSA)26Sor^tm14(CAG-tdTomato)Hze^/J [JAX 007914] (3 F, 4 M; also known as AI14), C57BL/6J-Tg(Thy1-GCaMP6s)GP4.12Dkim/J [JAX 026776] (8 F, 4 M; also known as Thy1-GCaMP6s), B6.Cg-Tg(Syn1-cre) 671Jxm/J [JAX 003966] (2 F, 3 M), and cross between AI14 and Thy1-GCaMP6s (7 F, 11 M). All mice were housed in a room with a reversed light cycle (12–12 h). Experiments were performed during the dark period.

### IBL Decision-making task

For details, refer to Appendix 2 and 3 in the previous paper^20^. Briefly, all mice were first implanted with a headbar, then underwent water-restriction and habituation following the IBL protocols, before the task training began^20^. In the IBL task, head-fixed mice turn a steering wheel using their forepaws to move a visual stimulus (Gabor patch) on the screen (**Figure 1a**). When the stimulus is on the right (35 degrees in the azimuth, 0 degrees elevation relative to the center of the mouse visual field), mice must turn the wheel leftward (counterclockwise) within 60 seconds from stimulus onset, and rightward for the stimulus on the left, to receive a water reward (∼3 μL). In the beginning of training, only high-contrast stimuli (100% and 50% contrast) are presented to facilitate learning the stimulus-choice. Once performance (i.e., correct choice rate) exceeds 80%, low-contrast stimuli (25%, 12.5%, 6.125%, and 0%) are gradually introduced. Although a subset of mice in our study performed the task with all contrast levels in later training sessions after reaching the learning criterion - 80% in high-contrast trials, we used only the high-contrast trials as a measure of task performance in all sessions. Each session lasted 49 ± 9.0 (mean ± s.d.) minutes and mice performed 427 ± 158.0 (mean ± s.d.) trials per session.

### Choice behavior models

To compute the degree of persevere and exploit strategy in the choice sequence of each mouse, we used three slightly different models, all of which produced similar results (**Figure S4**). The first model is described in the following equation:

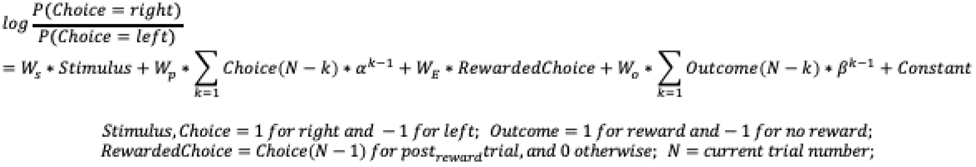

The first term corresponds to the current trial stimulus dependence. The second term represents perseverance as it captures the dependence on leaky accumulation of past choices. The third terms capture the ‘win-stay’ strategy (i.e., exploitation). The fourth term was added as our previous studies found many mice exhibit previous trial outcome dependence in ambiguous choice contexts^17,18^. We estimated the weights and constant that minimize the negative log likelihood of the observed choice sequence using a Python Scipy function *minimize*.

The second model replaced the exponential decay in choice and outcome accumulation with discrete weights for each n-back trial as follows:

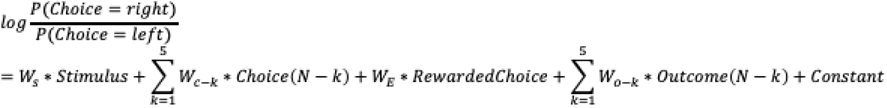

In the last model, we removed the outcome-dependence term and then added a term to capture the tendency to repeat trials following no reward as follows:

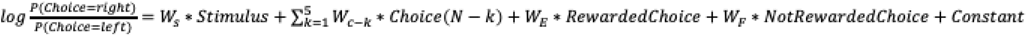

When assessing differences between LED-on and off trials in the optogenetic experiment, we doubled the number of weights in the first model so that one set is for LED-on trials, while the other for LED-off trials.

### Exploration score

Both the perseverance and exploitation weights in the model would decrease as exploratory choices are enhanced. Based on this inverse relationship, we computed a single exploration score by combining the two weights using the following formula:

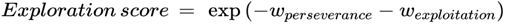

The exploration score becomes closer to 0 as perseverance and exploitation strengthen. In contrast, the exploration score becomes close to 1 when perseverance and exploitation decrease to 0, and exceeds 1 when those weights change their signs.

### Learning rate

To compute a learning rate, using a Python Scipy optimization function *curve_fit*, we fit the learning curve of each mouse with the following exponential function:

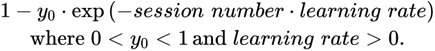

### Wheel movement kinematic analysis

A time series of wheel position data was first uniformly re-sampled at 1 kHz before we took the time-derivative of the resampled position data as velocity. We detected movement onset and offset using empirically determined threshold and duration (i.e., |velocity|>0.2 rad/s continuously for 10 ms for onset; |velocity|<0.1 rad/s continuously for 200 ms for offset). Each trial could contain multiple discrete movement bouts, each defined with distinct movement onset and offset. Response time was measured as time from stimulus onset to the movement onset of the last bout that led to feedback. Peak speed was the maximum amplitude of the velocity in the last bout. The trial-to-trial correlation was computed using the first 200-ms long velocity time series during the last bout.

### Optogenetic experiment

Mice (N=7; 4 F, 3 M; 6 wild type, 1 AI14::Thy1-GCaMP6S) were implanted with a headbar and then injected bilaterally in PPC (2.0 mm posterior to bregma, 1.7 mm lateral from the midline, and 0.5 mm beneath the dura) with a mixture of pENN.AAV.hSyn.Cre.WPRE.hGH (Addgene 105553-AAV1) and pAAV-nEF-Con/Foff2.0-ChRmine-oScarlet (Addgene 137161-AAV8) (100 nL each hemisphere). Approximately two weeks after injection, we implanted fiber optic cannulas (0.2 mm core, 0.37 NA, 1.25 mm stainless steel ferrule; Newdoon) into each side of PPC, at 0.5 mm beneath the dura through bur holes. Optogenetic experiment mice underwent the same water-restriction, habituation, and training procedures as the regular control mice, with the exception that in the first 15-16 sessions, their fiber optic cannulas were connected to the LED light source (625 nM, ∼ 10 mW; Thorlabs M625L4) to stimulate PPC during the ITI (4.0 ± 0.68 (mean ± s.d.) seconds) on ∼23% of randomly selected trials. Following the optogenetic sessions, mice continued daily training in 10-15 more sessions without stimulation.

### Statistical analysis

We used the Wilcoxon rank-sum test for comparing two groups of independent samples, Wilcoxon signed-rank test for paired samples, Kruskal-Wallis one-way ANOVA for comparing multiple groups of independent samples, Spearman rank correlation coefficient for assessing correlation between two groups of variables, and chi-square test for comparing categorical distributions between groups of independent samples. When comparing the performance between two nested models for choice behavior (full versus simple model), we used the likelihood ratio test.

## ACKNOWLEDGEMENT

We thank Dr. Bardia F. Behabadi for his technical support with implementing the IBL task and data analysis, and Terry Cirrencione, Ryan Wang, Elizabeth Croce, Eric Cho, Adam Schwartz, Keana Keen, Aylin Sanroman, and Kameron Kaplan for their help with animal training. This work was supported by grants to E.J.H. (NIH R56-MH130488, Sloan Research Fellowship, and Schweppe Scholar Award).

## AUTHOR INFORMATION

E.J.H. conceived the project, executed experiments, analyzed data, and wrote the paper. S.K., Y.H., and J.S. executed experiments, analyzed data, and wrote the paper. B.L. executed experiments and analyzed data. A.O., M.D., L.M.W., N.P., and S.B executed experiments.

