## Supplementary Figures for "Parietal stimulation reverses age-related decline in exploration, learning, and decision-making"

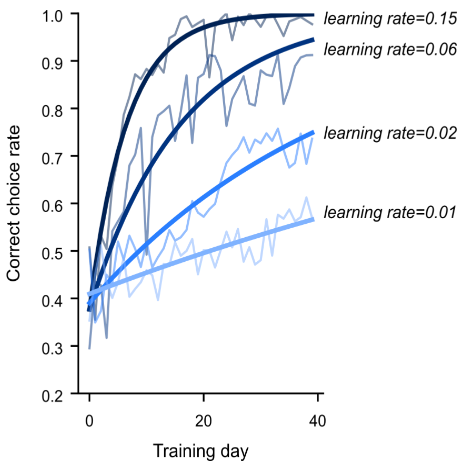


**Figure S1. Learning rate estimates from exponential function fit to learning curves.** Four example individual learning curves in different colors are fit with exponential functions. The time constant of each exponential function is used as the learning rate.

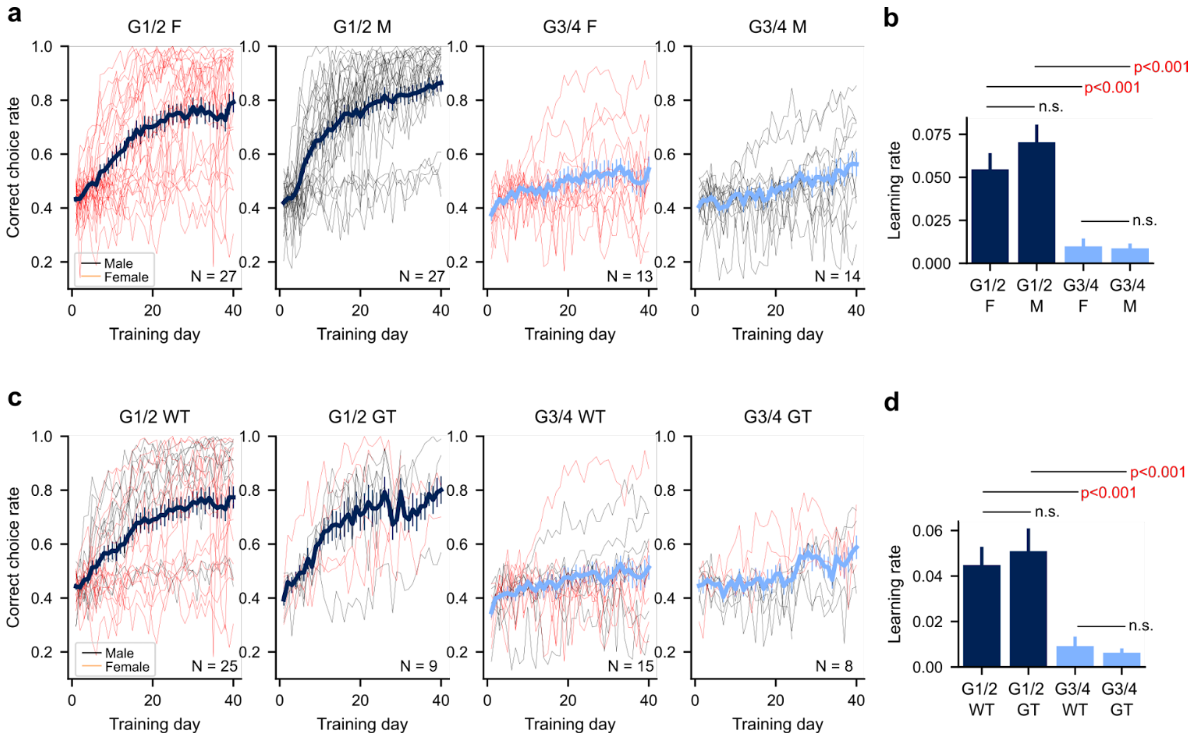


**Figure S2. Age-related learning declines are similar across sexes and strains.**

- **a.** G1 and G2 are combined as a young mouse group, while G3 and G4 are combined as an old mouse group. Learning curves of each age group from each sex separately. Thick lines: mean ± S.E.
- **b.** The learning rate of each age and sex group. Mean ± S.E.
- **c.** Learning curves from the two groups, separated for two different strains, wild type (WT) versus cross between Thy1-GCaMP6S and LSL-tdTomato (GT).
- **d.** The learning rate of each age and strain group.

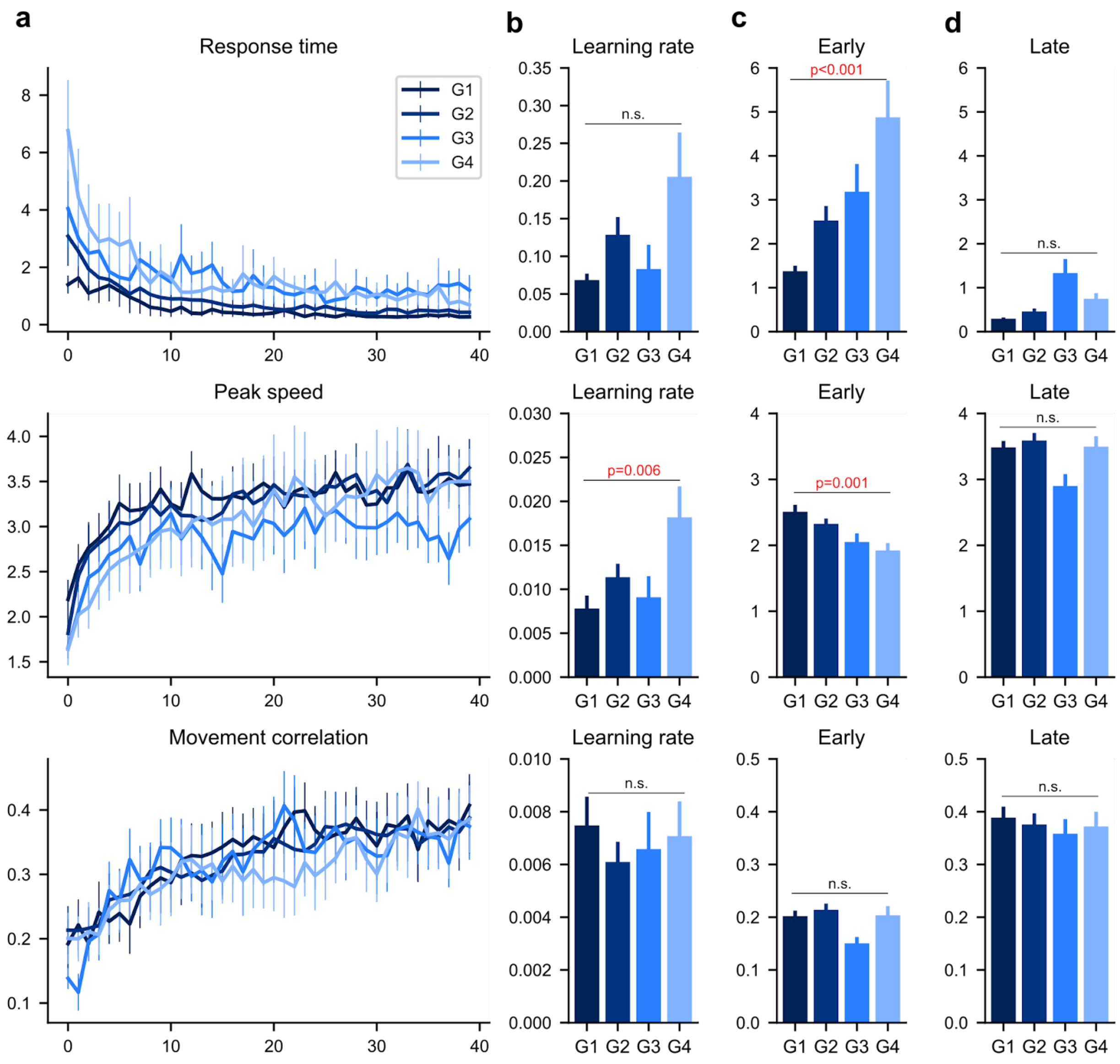


**Figure S3. Repetition-based motor learning is preserved with age.**

- **a.** Learning curves for three kinematic metrics, response time, peak movement speed, and trial-to-trial movement correlation. Mean ± S.E.
- **b.** The learning rate of the three kinematic variables across the four different age groups. Mean ± S.E.
- **c.** The mean values in the first three training days.
- **d.** The mean values in the last three training days.

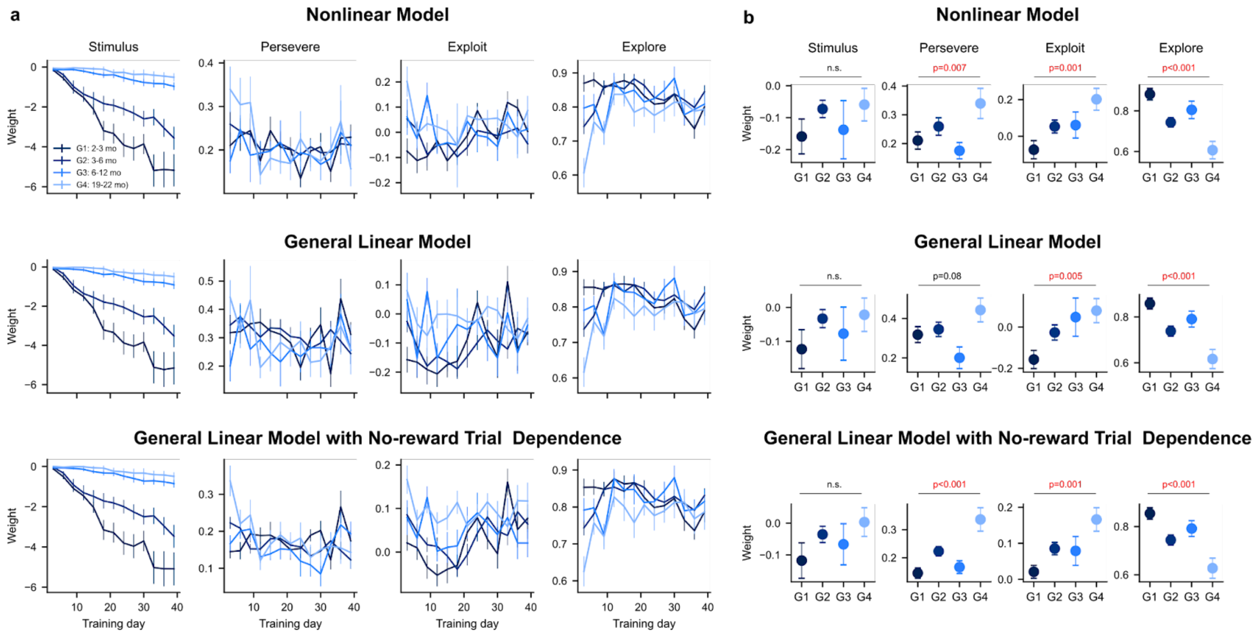


**Figure S4. Different variants of choice models produce similar results.**

**a.** Nonlinear history mode: the same model as in Figure 2c. Mean ± S.E.

**b.** General linear model: The weight of each N-back term in the choice-history and outcome history terms were independently estimated, rather than constraining them to exponentially decay with temporal distance from the current trial.

**c.** In this model, outcome history dependence was removed, but non-rewarded choice dependence was added.
